# B-Cell Precursor Acute Lymphoblastic Leukemia elicits an Interferon-α/β response in Bone Marrow-derived Mesenchymal Stroma

**DOI:** 10.1101/2023.05.09.539232

**Authors:** Mandy W. E. Smeets, Elisabeth M. P. Steeghs, Jan Orsel, Femke Stalpers, Myrthe M. P. Vermeeren, Christina H. J. Veltman, Stefan Nierkens, Cesca van de Ven, Monique L. den Boer

## Abstract

B-cell precursor acute lymphoblastic leukemia (BCP-ALL) can hijack the normal bone marrow microenvironment to create a leukemic niche which facilitates blast cell survival and promotes drug resistance. Bone marrow-derived mesenchymal stromal cells (MSCs) mimic this protective environment in *ex vivo* co-cultures with leukemic cells obtained from children with newly diagnosed BCP-ALL. We examined the potential mechanisms of this protection by RNA sequencing of flow-sorted MSCs after co-culture with BCP-ALL cells. Leukemic cells induced an interferon (IFN)-related gene signature in MSCs, which was partially dependent on cell-cell signaling by tunneling nanotubes. The signature was selectively induced by BCP-ALL cells, most profoundly by *ETV6-RUNX1* positive ALL cells, as co-culture of MSCs with healthy immune cells did not provoke a similar IFN signature. Leukemic cells and MSCs both secreted IFNα and IFNβ, but no IFNγ. In line, the IFN-gene signature was sensitive to blockade of IFNα/β signaling, but less to that of IFNγ. The viability of leukemic cells and level of resistance to three chemotherapeutic agents was not affected by interference with IFN signaling using selective IFNα/β inhibitors or silencing of IFN-related genes. Taken together, our data suggest that the leukemia-induced expression of IFNα/β-related genes by MSCs does not support survival of BCP-ALL cells but may serve a different role in the pathobiology of BCP-ALL.

## Introduction

B-cell precursor acute lymphoblastic leukemia (BCP-ALL) is the most common pediatric malignancy, and is characterized by malignant transformation and clonal expansion of B-precursor cells in the bone marrow ^1–3^. The 5-year event-free survival rate of pediatric patients with BCP-ALL is currently ∼90% ^4–8^. However, despite a relatively high cure rate, BCP-ALL still represents a major cause of cancer-related mortality in children, which can be attributed to treatment-related death or relapse of the leukemia ^6,9^.

Several studies have shown that the bone marrow microenvironment facilitates leukemogenesis and contributes to cellular drug resistance ^9–15^. Bone marrow-derived mesenchymal stromal cells (MSCs) are a major component of the bone marrow microenvironment ^16^. *Ex vivo*, these cells provide a survival benefit to co-cultured leukemic cells and induce resistance to chemotherapeutic drugs. We previously showed that leukemic cells re-sensitize to chemotherapeutics *ex vivo* when the interaction with MSCs is being disrupted, e.g., by interfering with tunneling nanotube (TNT) formation ^10^. Leukemic cells use TNTs as an effective mechanism to communicate with MSCs. Transfer of mitochondria, autophagosomes, and transmembrane proteins towards MSCs has been established ^14^. Disruption of TNTs resulted in an altered cytokine profile secreted by MSCs and a reduced survival benefit for leukemic cells ^10,14^. These findings indicate that MSCs play an important role in the maintenance of BCP-ALL, for which however the mechanism is yet unknown. To elucidate potential mechanisms, we investigated the effect of leukemic cells on the gene expression profile of MSCs. Subsequently, we studied whether differentially expressed genes contributed to the viability and drug responsiveness of BCP-ALL. Our study shows that the interferon (IFN) α/β pathway is selectively activated in MSCs upon interaction with BCP-ALL cells but not with normal, healthy immune cells. However, interference with this pathway did not affect the viability nor level of drug resistance, suggesting that the IFNα/β response may serve a different role in the pathobiology of BCP-ALL.

## Materials and Methods

### Mesenchymal stromal cells

Primary MSCs were obtained from pediatric bone marrow aspirates taken at diagnosis of BCP-ALL or at relapse. MSCs were selected based on adherence capacity, followed by confirmation of negativity for surface markers CD19/CD34/CD45, and positivity for CD44/CD54/STRO-1/CD73/CD90/CD105/CD146/CD166. MSCs were passaged twice a week in DMEM low glucose/pyruvate/HEPES medium (Gibco, Thermo Fisher Scientific, Waltham, MA, USA) supplemented with 15% fetal calf serum (FCS) (Bodinco BV, Alkmaar, The Netherlands), amphotericin B (1:166; 250μg/ml, Gibco), gentamycin (1:1000; 50ng/ml, Gibco), fresh 1ng/ml recombinant human fibroblast growth factor (FGF) basic (BioRad, Veenendaal, The Netherlands) and 0.1mM L-ascorbic acid (Sigma Aldrich, Zwijndrecht, The Netherlands). MSCs were cultured at 37°C/5%CO2 and used for experiments until passage 10. For this study, MSCs from three different donors were selected (***Supplementary table 1***).

### Primary BCP-ALL cells

Primary BCP-ALL cells were collected from bone marrow samples originating from children with newly diagnosed BCP-ALL (1-18 years). This study was performed in agreement with the Institutional Review Board, and written informed consent was given by patients, parents, or guardians. As described by Den Boer *et al*. ^17^, isolation and processing of the leukemic blasts was performed by density gradient centrifugation using Lymphoprep (1.077 g/ml, Nycomed Pharma, Oslo, Norway) for 15 minutes at 1500rpm. When necessary, normal hematopoietic cells were depleted using magnetic beads coupled to monoclonal antibodies to enrich for leukemic cells, resulting in ≥ 90% blasts for all samples prior to experiments. Primary BCP-ALL cells were kept in (short-term) culture at 37°C/5%CO2 using RPMI 1640 Dutch Modified medium (Gibco) containing 20% FCS (Bodinco), 2% PSF (penicillin 5.000U/ml, streptomycin 5.000U/ml, fungizone 250μg/ml, Gibco, Thermo Fisher Scientific), gentamycin (0.2mg/ml, Gibco), 1% insulin-transferrin-selenium (ITS) (Sigma Aldrich), and 2mM L-glutamin (Life Technologies, Thermo Fisher Scientific). A total of 15 primary BCP-ALL samples was selected for this study, including 8 *ETV6-RUNX1*, 1 high hyperdiploid, and 6 B-other cases (***Supplementary table 2***).

### Isolation and characterization of healthy immune cells from umbilical cord blood

Cord blood was collected by and received from the Vlietland Hospital in Schiedam. Mononuclear cells were isolated from umbilical cord blood using Lymphoprep (1.077 g/mL, Nycomed Pharma) as described above. The collection and use of umbilical cord blood from donors for our studies was approved by the ethics committee of the Erasmus University Medical Center and the scientific research committee of the Vlietland Hospital. Antibodies against B-cell, T-cell, NK cell, monocyte, granulocyte, and (plasmacytoid) dendritic cell markers were purchased for characterization of immune cells using flow cytometry: PerCP/Cy5.5 anti-human CD3 and CD11c (Biolegend, San Diego, CA, USA), PE anti-human CD4 and CD56 (Biolegend), Brilliant violet (BV)421 anti-human CD8 (Biolegend), Vioblue anti-human CD14 (Miltenyi Biotec), BV650 anti-human CD16 (Biolegend), BUV395 anti-human CD19 and CD123 (BD Biosciences, San Jose, CA, USA), BV605 anti-human CD45 (Biolegend), APC anti-human CD66b, and HLA-DR (Biolegend). Antibody panel 1 includes CD3/CD4/CD8/CD16/CD19/CD45/CD66b antibodies; antibody panel 2 includes CD11c/CD14/CD16/CD45/CD56/CD123/HLA-DR antibodies. ViaKrome808 Fixable Viability Dye (Beckman Coulter, Brea, CA, USA) was used to select for the viable population. Immune cells were characterized using the Cytoflex LX (Beckman Coulter). Flow cytometry data were analyzed using FlowJo (version 10.7.1).

### BCP-ALL cell lines

The cell lines REH (*ETV6-RUNX1*), Nalm6 (*DUX4-*rearranged B-other), RCH-ACV (*TCF3-PBX1*), SupB15 (*BCR-ABL1*), and MUTZ5 (*CRLF2-*rearranged) were cultured at 37°C/5%CO2 in RPMI 1640 GlutaMAX medium (Gibco) containing either 10 or 20% FCS (Bodinco) and 2% PSF (Gibco) and were passaged twice a week. Authentication of cell lines was confirmed regularly by short tandem repeat (STR) profiling by using the GenePrint 10 kit (Promega, Madison, WI, USA) on an ABI 3730 DNA analyzer (Applied Biosystems, Waltham, MA, USA).

### Co-culture of BCP-ALL cells, healthy immune cells and MSCs

MSCs (0.225×10^6^) were seeded in a 6-well plate in 3 ml DMEM medium (15% FCS, gentamicin, FGF, and L-ascorbic acid), and cultured for 24-26 hours at 37°C/5%CO2. The MSC monolayer (∼95% confluent) was washed with PBS, followed by adding 2 ml RPMI-1640 Dutch Modified medium. A volume of 1 ml RPMI 1640 Dutch Modified medium containing 5.0×10^6^ primary BCP-ALL cells or healthy immune cells was added to the MSCs. In case of BCP-ALL cell lines, 1 ml RPMI 1640 medium containing 0.9×10^6^ cells was added. Mono-cultures of BCP-ALL cells and MSCs were used as controls. The MSCs and BCP-ALL cells were cultured for 40 hours at 37°C/5%CO2.

### Fluorescent activated cell sorting

Cells were harvested and stained with antibodies against B-cell, MSC and hematopoietic cell markers. Antibodies used to distinguish the different cell populations included Brilliant violet 421 anti-human CD19 (Biolegend, San Diego, CA, USA), PE anti-human CD45 (Biolegend), Alexa Fluor 750-conjugated human 5’-Nucleotidase/CD73, MCAM/CD146, and ALCAM/CD166 (R&D Systems, Minneapolis, MN, USA). Sytox Red Dead Cell Stain (Thermo Fisher Scientific) was used to select for the viable population. Viable cells were sorted by the SH800S Cell Sorter (Sony, San Jose, CA, USA) with a 100μm nozzle into an MSC- and leukemic cell fraction. A purity check of both fractions was performed by measuring 10,000 events after sorting (***Supplementary table 3***). The maximum percent of contaminating cells allowed in sorted fractions was determined by the effect these contaminating cells had on the expression levels of genes. This was previously determined in our lab by performing a quantitative RT-PCR experiment in which increasing amounts of MSCs, or ALL cells were added to ALL cells or MSCs, resp. In this study, we aimed for a maximum of 5% ALL cells in sorted MSC fractions, and only 0.3% MSCs in sorted ALL fractions as MSCs contain more RNA than ALL (***Supplementary Figure 1***). 17/45 (37.8%) sorted ALL samples contained >0.3% MSCs, while only 5/45 (11.1%) sorted MSC samples contained >5.0% ALL cells (***Supplementary Figure 1/Supplementary table 3)***. Although some samples showed high contamination, all samples were included for further analysis. The sorted MSCs and leukemic cells were pelleted, and dissolved in 1ml Trizol (Invitrogen, Bleiswijk, The Netherlands) for RNA isolation. Samples were stored at -80ºC until further use.

### Multiplexed fluorescent bead-based immunoassay (Luminex)

Supernatants of mono- and co-culture experiments were centrifuged for 10 minutes at 1200rpm to remove cellular debris. Supernatants were immediately stored at -80°C. An in-house developed and validated multiplex immunoassay based on Luminex technology (xMAP, Luminex, Austin, TX, USA) was used to measure cytokines of interest at the Center for Translational Immunology (UMC Utrecht, the Netherlands) ^18^. Archived supernatants were incubated with antibody-conjugated MagPlex microspheres for 1 hour at room temperature with constant shaking. Samples were incubated 1 hour with biotinylated detecting antibodies, followed by 10 minutes of incubation with phycoerythrin (PE)-conjugated streptavidin. PE-conjugated streptavidin was diluted in high performance ELISA buffer (HPE) (Sanquin, Amsterdam, The Netherlands). The Biorad FlexMAP3D (Biorad laboratories, Hercules, CA, USA) and xPONENT software version 4.2 (Luminex) were used to determine the concentration of the secreted cyto- and chemokines in supernatants. Bio-Plex Manager software, version 6.1.1 (BioRad) was used to analyze and to quantify the data using standard dilution series.

### RNA sequencing

RNA extraction was performed with Trizol (Invitrogen) as described by manufacturer. The quantity, and A260/A230 and A260/280 ratios of RNA were determined by the Qubit 4 fluorometer (Thermo Fisher Scientific), and the DeNovix Spectrophotometer DS 11 (DeNovix Inc., Wilmington, DE, USA), respectively. RNA quality was determined using the Bioanalyzer Agilent 2100 (Agilent, Santa Clara, CA, USA) combined with the Agilent RNA 6000 Nano Kit according to manufacturer’s protocol. 600ng RNA (25 ng/ul; RIN>6.8) was used for ribonucleotide-depleted long-noncoding RNA sequencing on a NovaSeq 6000 (Illumina, San Diego, CA, USA). Fastq reads were aligned to the human reference genome GRCh38 using STAR (version 2.6.0c) and data were analyzed using R (version 3.6.3). Genes with zero counts across all samples, or <1.0 counts per million mapped reads in all but one sample were removed. Normalization factors were calculated using the trimmed mean of M values (TMM) method. Dispersion in the data was estimated taking batch effect and subtype of BCP-ALL cells into consideration. EdgeR (version 3.28.1) was used to perform differential gene expression analysis. A paired EdgeR p-value corrected for multiple testing (FDR) below 0.05 was considered significant. Fragments per kilobase per million reads mapped (FPKM) values were generated using the transcript length of the longest known transcript (Gencode v29). FPKM values were log10-transformed to create a heatmap. Pathway analysis was performed using Pathvisio software ^19^.

### Tunneling nanotube inhibition

Vybrant DiI Cell-Labeling Solution (Invitrogen) was used to pre-stain BCP-ALL cell line REH for 24 hours. Transwells (0.4μm pore-size; Corning, New York, USA) were pre-incubated with RPMI medium for 30 minutes at 37°C/5%CO2 before DiI-stained ALL cells were added. Dye transfer from DiI-stained ALL cells cultured in transwells towards MSCs cultured in 6-well plates (Sigma Aldrich) was determined after 40 hours by flow cytometry, visualized by excitation with 561nm laser and emission detection with 585/42 filter using the Cytoflex S (Beckman Coulter). MSCs cultured with DiI-labeled ALL cells without transwell were used as control. Flow cytometry data were analyzed by using FlowJo (version 10.7.1) and graphs were created by Graphpad Prism (version 9.1.2).

### Lentiviral silencing

pLKO.1-puro Mission vectors (Sigma Aldrich), containing short-hairpin RNA (shRNA), were purchased to target genes involved in the Interferon (IFN) pathway. The pLKO.1-puro Mission vectors non-mammalian shRNA control plasmid and luciferase shRNA control plasmid (SHC002 and SHC007, resp.) were used as control vectors. Human embryonic kidney 293T (HEK293T) cells were cultured in DMEM, high glucose, glutamax medium (Gibco) supplemented with 2% PSF (Gibco) and 10% FCS (Bodinco). HEK293T cells were transfected at 80% confluence according to manufacturer’s protocol. In short, transfection was performed with 1.6μg pVSV-G (Addgene plasmid 12259; Addgene, Cambridge, Massachusetts, USA), 3.7μg pPAX2 (Addgene plasmid 12260), and 10.7μg plasmid DNA (Sigma Aldrich; *MX1* (#3: TRCN0000056880; #5: TRCN0000056882), *IFI6* (#4: TRCN0000057501), *IFI27* (#2: TRCN0000115858), *IFI44L* (#1: TRCN0000167757; #3: TRCN0000242541) or *ISG15* (#4: TRCN0000007423)) in the presence of X-tremeGENE HP DNA Transfection Reagent (Sigma Aldrich). Five shRNAs per gene were tested for knockdown efficiency, but merely shRNAs resulting in efficient knockdown (>60%) were used for consecutive experiments. Fresh medium was added 24 hours after transfection. Virus supernatant was harvested two and three days after transfection of HEK293T cells and filtered with a 0.45μm filter to eliminate cell debris. Virus particles were collected by centrifugation at 32,000rpm for 2 hours at 4°C using the Optima XE-90 ultracentrifuge (Beckman Coulter). Virus was aliquoted and stored at -80°C until further use. Virus titration was performed on primary MSCs to determine the optimal virus concentration. Genes of interest were silenced (KD) by transfecting MSCs using spin-infection (1800rpm, 45 minutes, 21°C), followed by selection for puromycin resistance for 1 week. RNA was extracted by means of the RNeasy Mini Kit (Qiagen, Hilden, Germany), and was used to determine the knockdown efficiency by quantitative RT-PCR.

Next, lentiviral transduced MSCs were seeded in 24-well plates (50,000 cells/well) and incubated for 5 hours at 37°C/5%CO2 in 700μl RPMI-1640 Dutch Modified medium. A volume of 175μl RPMI 1640 medium with 875,000 primary BCP-ALL cells was added to the MSCs (1.0×10^6^ BCP-ALL cells/ml). Mono- and co-cultures were performed for 72 hours, after which viability of MSCs and BCP-ALL cells was determined by flow cytometry (Cytoflex S) with the antibodies described previously.

### Activation and inhibition of interferon signaling

MSCs were stimulated with recombinant human IFNα (0.03 U/ml) or IFNβ (60 pg/ml) protein (R&D Systems), in combination with different concentrations of inhibitor of IFNα/β (i-IFNα/β) (Recombinant Viral B18R, R&D Systems; 0.9, 1.8 or 3.6 ng/ml) for 40 hours. Inhibitors of IFNα/β (1.8 ng/ml), and IFNγ (Recombinant Viral B8R, R&D Systems; 3.0 ng/ml), by functioning as decoy receptors, were used in unstimulated co-cultures of MSCs and BCP-ALL cells for 40 hours. Cell viability and the IFN-related gene expression signature were assessed with flow cytometry and quantitative RT-PCR, resp.

### Drug exposure experiments

BCP-ALL cells in mono- or co-culture with MSCs were exposed to a range of six concentrations of L-asparaginase (ASP, 0.003 – 10 IU/ml), daunorubicin (DNR, 0.002 – 2 μg/ml) or prednisolone (PRED, 0.008 – 250 μg/ml) to create a dose-response curve. MSCs (5,500) were seeded in 96-well flat-bottom plates and incubated for 24 hours. Primary BCP-ALL cells (137,500; 1.0×10^6^/ml) were added with or without ASP, DNR or PRED, and were incubated for 4 days at 37°C/5%CO2. Cells were harvested, stained with the antibodies described previously, and the number and percentage of viable Sytox Red negative cells was measured by means of flow cytometry (Cytoflex S). Sensitivity of leukemic cells to L-asparaginase (ASP), daunorubicin (DNR) and prednisolone (PRED) in mono- and MSC co-culture in the presence of i-IFNα/β was tested. The concentration of drug most discriminative for resistance induced by MSCs was determined per BCP-ALL sample (***Supplementary Figure 2***). These drug concentrations were tested with and without i-IFNα/β (1.8 ng/ml). In detail, 16,500 MSCs were seeded in 48-well plates, incubated for 24 hours, followed by addition of 412,500 primary leukemic cells (1.0×10^6^/ml) with or without i-IFNα/β. The previously determined optimal drug concentration was added to the cells and incubated for 4 days at 37°C/5%CO2. Cells were harvested and measured using flow cytometry.

### Quantitative RT-PCR (RT-qPCR)

Primer-probe pairs were purchased at Thermo Fisher Scientific targeting IFN-related genes. These genes included *MX1* (Hs00895608_m1), *IFI6* (Hs00242571_m1), *IFI27* (Hs01086370_m1), *ISG15* (Hs00192713_m1), *IFI44L* (Hs00915292_m1), *CXCL10* (Hs00171042_m1), *IFIT1* (Hs01675197_m1), *OAS3* (Hs00196324_m1), *IFITM1* (Hs00705137_s1), and housekeeping gene *GAPDH* (Hs99999905_m1). RNA was extracted using the RNeasy Mini Kit (Qiagen) according to manufacturer’s protocol. cDNA was synthesized by using the SensiFAST cDNA Synthesis kit (Bioline, London, England). Gene expression levels were measured using a MicroAmp Fast Optical 96-well Reaction Plate with Barcode, 0.1 ml (Thermo Fisher Scientific), and a QuantStudio 12K Flex Real-Time PCR System (Thermo Fisher Scientific). mRNA expression values were corrected for GAPDH (ΔCt = Ct_gene_ -Ct_GAPDH_). Fold change in relative gene expression was calculated using the 2^^-ΔΔ^Ct method. Graphs were made using GraphPad Prism Version 9.1.2. The IFN-related gene signature in MSCs upon (in)direct co-culture was measured by means of quantitative RT-qPCR on flow-sorted cells. mRNA expression levels in MSCs upon co-culture were normalized to MSC mono-culture (100%).

### Statistical analysis

One-way ANOVA tests and one-sided (un)paired t-tests were performed using Graphpad Prism, and correction for multiple testing was included when appropriate. A p-value below 0.05 was considered statistically significant: * p < 0.05, ** p < 0.01, *** p < 0.001, **** p < 0.0001.

## Results

### BCP-ALL induces an Interferon-related gene signature in MSCs

RNA sequencing was performed on sorted fractions of MSCs after exposure to primary BCP-ALL patient samples (n=15; **Figure 1A**). Pathway analysis of RNA sequencing data revealed enrichment for IFN-pathway activated genes in the MSCs after co-culture with BCP-ALL cells (***Supplementary Figure 3***). All three tested sources of bone marrow-derived MSCs (MSC#1-3) increased the IFN-related gene expression levels in co-culture with BCP-ALL cells, suggesting that the leukemia-induced change in expression is independent from the source of MSCs (**Figure 1B**). The IFN-related genes including *IFI6, MX1, IFI27*, and *OAS3* were 4.3 to 7.7-fold upregulated in MSCs upon co-culture with BCP-ALL compared to MSCs kept in mono-culture (FDR=1.08×10^−16^, FDR=2.23×10^−15^, FDR=3.81×10^−11^, FDR=1.44×10^−15^, resp.) (**Figure 1B**). Other IFN-related genes such as *IFITM1, IFI44L, IFIT1* and *ISG15* were 3.1 to 4.3-fold upregulated (FDR=1.08×10^−6^, FDR=5.21×10^−11^, FDR=2.99×10^−12^, and FDR=4.73×10^−12^ resp.; ***Supplementary Figure 4***). Expression of IFN-related genes was remarkably higher in MSCs after co-culture with *ETV6-RUNX1* BCP-ALL (n=8) compared to MSCs sorted after co-culture with B-other BCP-ALL cells (n=6) as exemplified for *IFI6* 3-fold, p=0.031, *MX1* 2.6-fold, p=0.048 and *OAS3* 2.1-fold p=0.04 in **Figure 1C** and for a set of 20 IFN signature genes in **Figure 1D**. Similar to our observation in primary patients’ samples (**Figure 1C, 1D**), the IFN-related profile was most strongly induced by the *ETV6-RUNX1* positive cell line REH but less by cell lines with another BCP-ALL subtype (***Supplementary Figure 5***). Although we did observe induction of *CXCL10* expression, no complete IFN signature in MSCs was induced upon exposure to healthy immune cells from cord blood (n=5) (**Figure 2A-C**).

**Figure 1.**
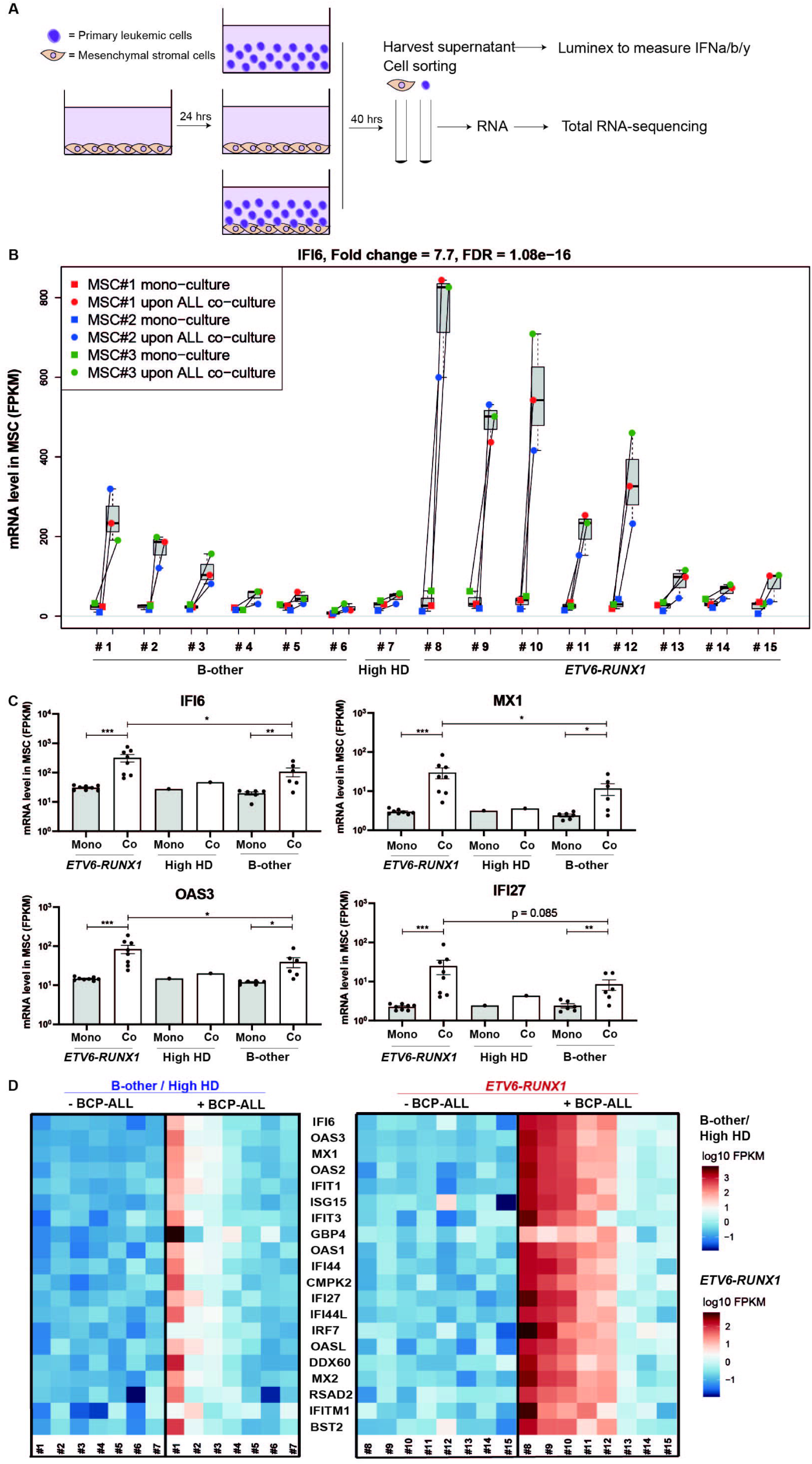
BCP-ALL cells induce Interferon-related gene signature in MSCs. (A) Flow chart of experimental setup. RNA sequencing was performed on sorted viable MSCs fractions after mono-culture and after MSC/ALL co-culture, for gating strategy see *Supplementary Figure 1*. Secreted interferon levels were measured in equal volumes of supernatant of mono-cultures and MSC/ALL co-cultures (Figure 6). (B) *IFI6* mRNA levels (FPKM) expressed in paired samples of MSCs sorted after mono-culture (squares; MSC#1-3 indicated in red, blue, and green, resp.) and MSCs sorted after co-culture with BCP-ALL cells from 15 individual patients (#1-15; circles). Boxplots represent the interquartile range, the median is depicted by a line. FDR, p-value false discovery rate. (C) mRNA expression levels in MSCs of *IFI6, MX1, OAS3* and *IFI27* upon mono-culture (grey bars) and co-culture (white bars) summarized per BCP-ALL subtype (n=8 for *ETV6-RUNX1*, n=1 for high hyperdiploid, n=6 for B-other). Each dot represents the average expression of IFN gene in 3 MSCs per sort. Mean ± SEM is indicated. (D) Heatmap of expression levels of IFN-signature genes in MSCs sorted after mono-culture (-ALL) and those after co-culture with 15 different ALL cases (+ALL). Heatmap is grouped by BCP-ALL subtype: high hyperdiploid (HD) and B-other ALL cases (#1-7) or *ETV6-RUNX1* ALL cases (#8-15). Blue, low expression. Red, high expression.

**Figure 2.**
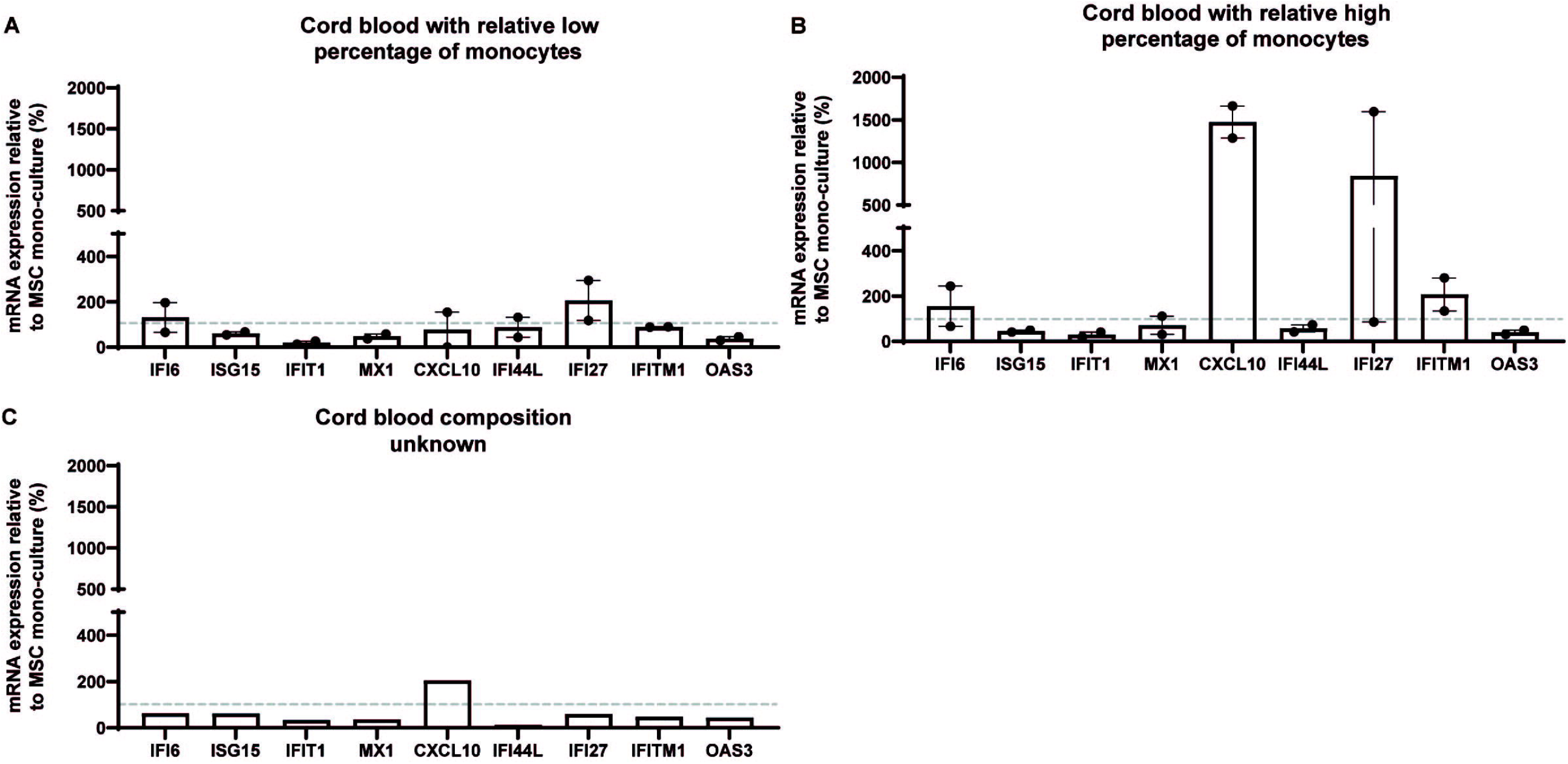
Interferon-related genes in MSCs are specifically induced by leukemic cells. *IFI6, ISG15, IFIT1, MX1, CXCL10, IFI44L, IFI27, IFITM1* and *OAS3* gene expression levels measured by RT-qPCR in MSCs (MSC#2) sorted after co-culture with mononuclear cells from five different cord blood samples: cord blood with relative (A) low (n=2) or (B) high (n=2) percentage of monocytes (>20%), or (C) unknown cellular composition (n=1). Bars are means of triplicate measurements ± SEM for 2 independent co-culture experiments. Dashed line (---) indicates mRNA expression levels of MSC after mono-culture, set at 100%.

In previous studies we showed that the viability of leukemic cells depends on a close and direct contact with MSCs mediated via tunneling nanotubes (TNTs) ^10^, as can be visualized by reduced DiI dye transfer from pre-labeled ALL cells in the upper compartment to MSCs in the bottom compartment of a transwell setting (**Figure 3A-B**; 87.6% decrease (p < 0.0001)). Although preventing the formation of TNTs significantly reduced the expression levels of *CXCL10* (3.3-fold, p=0.016), *IFI44L* (1.57-fold, p=0.009) and *IFITM1* (1.5-fold, p=0.005), not all tested genes were affected (i.e., *IFI6, IFITM1, MX1, IFI27*; **Figure 3C**). This suggests that TNT-dependent signaling partly contributes to the BCP-ALL induced gene expression changes in MSCs.

**Figure 3.**
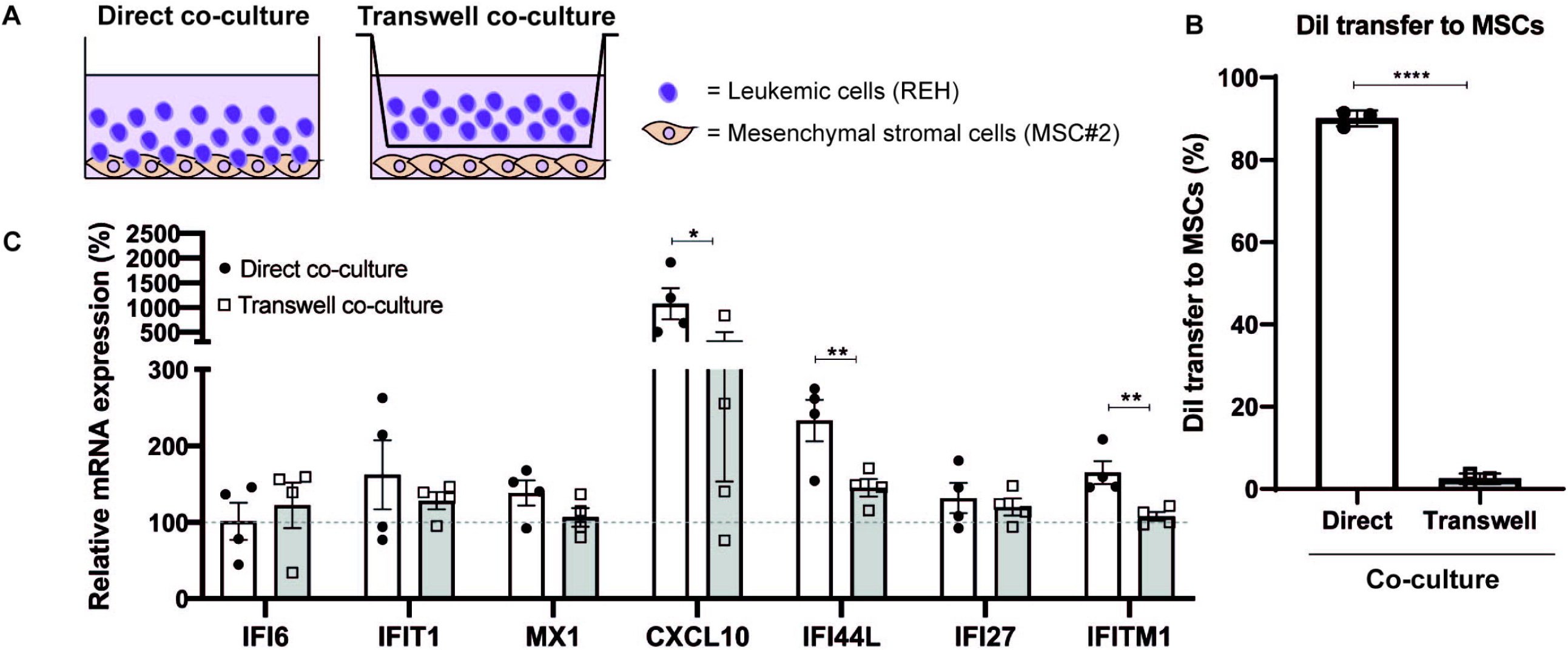
Direct cell-cell contact contributes to induction of Interferon-related gene expression in MSCs. (A) Schematic overview of experimental setup of direct co-culture and transwell co-culture of REH cells with MSC#2 using 0.4μm pore-size transwells (B) Percentage of DiI positive MSCs after direct (circles, white bars) and transwell (squares, grey bars) co-culture with REH leukemic cells for 40 hours. Dots represent averaged duplicate measurements; bars represent means ± SEM for 3 independent experiments. (C) IFN-related gene expression levels, measured by RT-qPCR, in MSCs sorted after direct (circles, white bars) or transwell (squares, grey bars) co-culture with REH cells for 40 hours. Dots represent averaged triplicate measurements; bars represent means ± SEM for 4 independent experiments. Dashed line (---) indicates mRNA expression levels of MSC after mono-culture, set to 100%.

To examine whether the upregulation of IFN-related genes in MSCs causally affected the viability of primary BCP-ALL cells, *IFI6, MX1, IFI27, IFI44L* and *ISG15* were lentivirally silenced resulting in an efficient knockdown of 67-93% for individual genes (***Supplementary Figure 6***). Efficient knockdown of *CXCL10* in MSCs could not be achieved (data not shown). Silencing of the strong IFN gene signature observed in MSCs upon exposure to *ETV6-RUNX1*-positive cells did not markedly decrease the viability of primary *ETV6-RUNX1* (nor B-other) BCP-ALL cells in 3 days culture assays (**Figure 4**).

**Figure 4.**
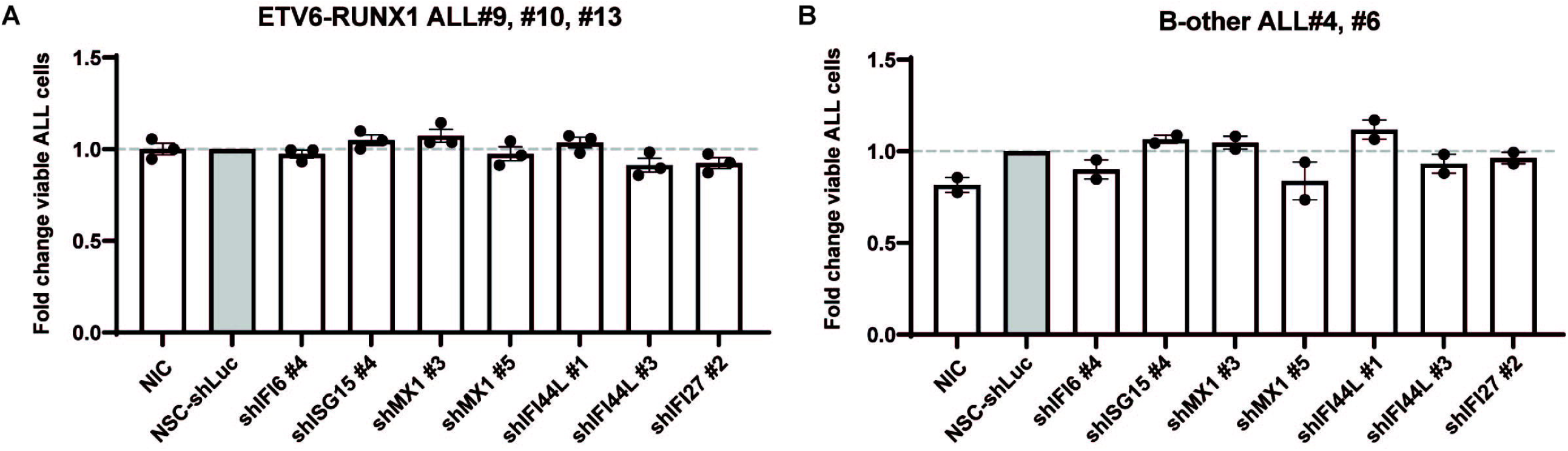
Silencing of Interferon-related genes in MSCs does not affect the viability of primary BCP-ALL cells. Fold change in percentage of viable leukemic cells after 72 hours of co-culture with transduced MSCs (MSC#2). Non-Silencing Control -short-hairpin-Luciferase (NSC-shLuc) is used as reference. (A) *ETV6-RUNX1* (n=3), and (B) B-other (n=2) BCP-ALL cells. Five shRNAs per gene were tested for knockdown efficiency; only shRNAs resulting in >60% knockdown (shIFI6 #4, shISG15 #4, shMX1 #3/5, shIFI44L #1/3, shIFI27 #2, see materials and methods) were used. Bars represent means ± SEM of duplicate measurements for 3 independent experiments. Dashed line (---) indicates that viability of ALL cells is equal to viability of ALL cells co-cultured with NSC-shLuc-MSCs (fold-change = 1; grey bars). NIC = Non-Infected Control.

### BCP-ALL triggers an IFNα/β but not IFNγ response in MSCs

BCP-ALL cells and MSCs both secreted quantifiable levels of IFNα and IFNβ, whereas secreted levels of IFNγ were below level of quantifiable detection (**Figure 5**). These secreted levels did not change upon co-culture of BCP-ALL and MSCs. Addition of recombinant IFNα and IFNβ to mono-cultures of MSCs clearly induced the expression of the IFN-index gene *IFI6*. This upregulation was prevented upon simultaneous addition of inhibitors of IFN signaling (i-IFNα/β) resulting in a 59-fold and 27-fold reduction in the IFNα and IFNβ induced expression of *IFI6* (**Figure 6A**). The BCP-ALL induced expression of IFN-related genes in co-cultures of MSCs was also reduced upon exposure to i-IFNα/β but not to i-IFNγ (**Figure 6B**). The survival benefit for ALL cells induced by MSCs (1.84-fold, p=0.003, **Figure 6C**) was not sensitive to IFNα/β inhibition (**Figure 6C**). Inhibitors of IFNα/β did neither sensitize BCP-ALL cells to L-asparaginase, daunorubicin and prednisolone (**Figure 6D**).

**Figure 5.**
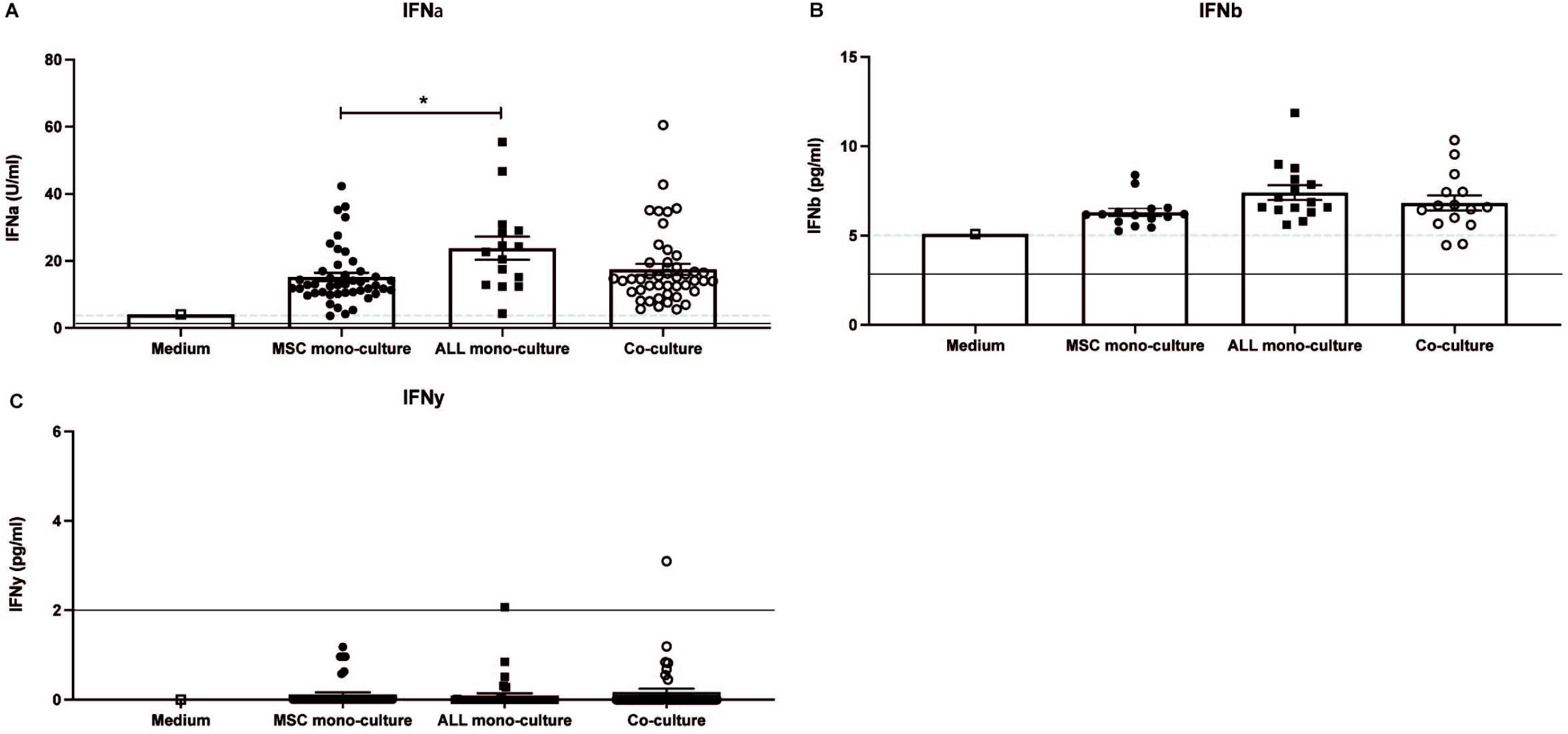
IFNα and IFNβ but not IFNγ is secreted by MSCs and primary BCP-ALL cells. IFN concentrations in culture medium and supernatant from 15 MSC and BCP-ALL collected after 40 hours of mono- and co-cultures. The concentration of IFN-cytokines is depicted, including (A) IFNα (U/ml), (B) IFNβ (pg/ml) and (C) IFNγ (pg/ml). Means ± SEM are shown. Dashed line (---) indicates concentration of IFNα, IFNβ or IFNγ present in culture medium. Limit of detection is indicated by solid line. Range of detection: IFNα: 0.5 – 2503 pg/ml, IFNβ: 2.8 – 14.926 U/ml, IFNγ: 2.0 – 12.342 pg/ml. IFNγ levels were below the level of linear detection for all but two samples.

**Figure 6:**
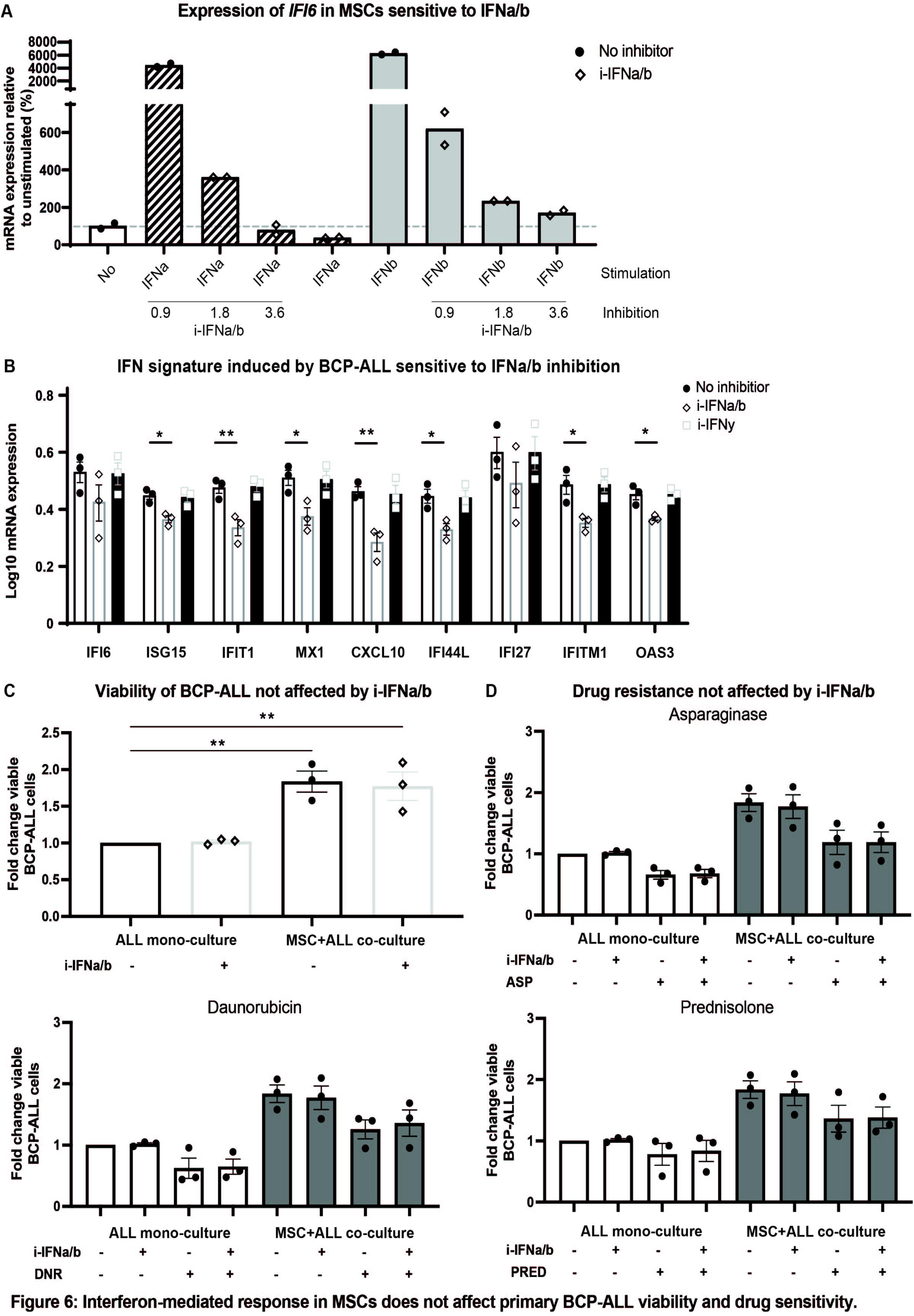
Interferon-mediated response in MSCs does not affect primary BCP-ALL viability and drug sensitivity. (A) Representing experiment showing technical replicates of RT-qPCR measured mRNA expression levels of *IFI6* in MSCs (MSC#2) upon 40 hours culture with recombinant human IFNα (striped bars), or -β (grey bars) protein (3.0U/ml and 60pg/ml, resp.) and different concentrations of i-IFNα/β (0.9, 1.8 or 3.6ng/ml) (diamonds), relative to unstimulated MSCs (white bar; circles). Dashed line (---) indicates mRNA expression levels of MSCs after mono-culture, set to 100%. (B) Log10-transformed IFN-related gene expression levels, measured by RT-qPCR, in unstimulated MSCs after 40 hour co-culture with primary BCP-ALL cells (ALL#9, #10, and #11) with or without IFNα/β (1.8ng/ml; diamonds) or IFNγ (3ng/ml; squares) inhibitors relative to MSC co-culture without i-IFN (circles). Bars represent means ± SEM of triplicate measurements for 3 independent experiments. (C) Fold change viable BCP-ALL cell percentage upon mono-culture with i-IFNα/β (1.8ng/ml), MSC co-culture without i-IFNα/β, or MSC co-culture with i-IFNα/β (1.8ng/ml), normalized to mono-culture without i-IFNα/β. Bars represent means ± SEM of triplicate measurements for 3 independent experiments. (D) Fold change viable BCP-ALL cell percentage in mono-culture or MSC co-culture upon exposure to ASP, DNR, PRED (concentration as determined in *Supplementary Figure 2*) and/or i-IFNα/β (1.8ng/ml), normalized to mono-culture without drug exposure or i-IFNα/β. Bars represent means ± SEM of triplicate measurements for 3 independent experiments.

## Discussion

Leukemic cells communicate with components of the bone marrow microenvironment in such a way that protection against chemotherapy and survival of leukemic cells is stimulated ^2,9–13^. In this study, we showed that BCP-ALL cells induce an IFN-related gene signature in MSCs, which was dependent on IFNα/β signaling, but independent of IFNγ. IFNα and IFNβ (but not IFNγ) were secreted by both BCP-ALL cells and MSCs but these levels were not increased upon co-culture of both cell types. The effect of normal cord blood cells (including e.g., T-cells, monocytes, NK- and dendritic cells) on expression levels of these genes in MSCs was limited, strengthening the leukemia-driven origin of this IFN signature. BCP-ALL samples with an *ETV6-RUNX1* translocation were the most potent inducers of the IFN-related gene expression signature in MSCs. This accounted for both primary BCP-ALL samples as well as an *ETV6-RUNX1* cell line. Genetic alterations of leukemic cells may be directly linked to interactions between leukemic cells and its microenvironment ^2^. Dander *et al*. has shown *in vitro* that the *ETV6-RUNX1* fusion can trigger modifications in adhesion molecule expression and adherence capacity of B-precursor cells ^20^. The role of cell-cell contact is also illustrated by our observation that inhibition of TNT formation negatively affected the expression of several IFN responsive genes.

Members of the IFN family are known to combat viral infections, modulate immune responses, and stimulate antitumor activities ^21–23^. On the other hand, an opposing, pro-tumorigenic role for interferons has been described as well ^22,24,25^. Recently, interferons were shown to elicit immune suppressive mechanisms in cancer, which may promote cancer progression and induce therapy resistance ^24,25^. Several studies have investigated the expression profile of IFN-related genes in cells derived from distinct cancer types ^26–29^. A subset of IFN-related genes, including the genes we found upregulated in MSCs, i.e. *IFI27, OAS1, OAS3, MX1* and *ISG15*, was persistently overexpressed in tumor cells resistant to chemo- or radiotherapy ^26,27,29^. Unfortunately, we could not determine whether the IFN-related genes were also upregulated in leukemic cells. This was due to technical limitations in the flow-sorting procedure resulting in a low percentage of MSCs in ALL fractions. This cross-contamination mainly affected gene expression levels of ALL sorted fractions because MSCs contain more RNA than ALL cells. Contamination of leukemic cells in the sorted MSC-fraction did not affect IFN expression levels in MSCs as basal expression levels in ALL cells did not exceed the levels measured in MSCs (***Supplementary Figure 1E***). For future studies, single cell sequencing is the preferred choice to detect which genes become upregulated in leukemic cells after contact with MSCs. Our study clearly demonstrates that blockade of IFNα/β signaling did not counteract the level of MSC-induced viability benefit and drug resistance of BCP-ALL cells. Our data imply that the BCP-ALL induced IFNα/β signature in MSCs serves a different role, e.g., attracting other cells to the leukemic niche. Since IFNα/β are involved in regulating the innate immune system ^30–33^, (in contrast to the IFNγ role in regulating T-cell function), BCP-ALL cells may benefit from an environment in which more innate immune cell types are present. Interferons α/β are involved in recruitment, function, maturation and/or activation of immune cells, such as natural killer (NK) cells, monocytes and dendritic cells (DCs) ^30,33^. Cyto-/chemokine production (e.g. CCL2-5 and CXCL10) can be induced by IFNα/β and may help to recruit inflammatory monocytes and NK cells ^24,33^. Immune cell infiltration into the tumor microenvironment has been reported in several cancer types, thereby creating a rich environment of growth factors and cytokines ^34^. Constant production of IFNα/β can suppress immune cell functions, leading to a decrease of immunosurveillance. For example, a prolonged IFNα/β gene signature is associated with less proliferation and differentiation of DCs and suppression of NK cells ^33^. In solid tumors, CXCL10 was shown to promote the migration of NK cells towards the tumor, and once trapped in the tumor microenvironment these NK cells downregulated NKG2D, the receptor that is important in the defense against cancer by the immune system ^35^. This network of trapped, but not functional (innate) immune cells, resulted in a condition promoting tumor cell proliferation, survival, and metastasis ^34,35^. In line with this observation, the increased CXCL10 levels observed in plasma of newly diagnosed ALL patients (Smeets *et al*., manuscript submitted) and the ALL-induced expression of *CXCL10* in MSCs (Figure 3) may be indicative for a similar process of impaired function of normal immune cells occurring in the bone marrow of leukemia patients, as was recently suggested by Pastorczak *et al*. ^2^.

In conclusion, our study reveals that IFNα/β but not IFN-γ contribute to the IFN signature elicited by BCP-ALL in MSCs. The induction of IFN-related genes in MSCs did not affect the viability and level of drug resistance of BCP-ALL cells observed upon exposure to MSCs. Our data warrant further studies into the role of the BCP-ALL-induced IFNα/β response in chemotaxis of healthy immune cells to create a beneficial environment for BCP-ALL cells. Interestingly, the IFNα/β gene signature in MSCs was predominantly induced by BCP-ALL cells harboring the *ETV6-RUNX1* fusion. Although *ETV6-RUNX1* leukemia has a favorable prognosis ^36,37^, it is one of the most commonly occurring genetic subtypes in pediatric leukemia ^20,36,38^. For this subtype, but also less favorable subtypes of ALL, it is important to explore the role of IFNα/β induced genes since this may point to alternative, maybe less intensive ways to eradicate leukemic cells.

## Acknowledgements

We would like to thank all group members of the Erasmus Medical Center in Rotterdam, and Princess Máxima Center in Utrecht for their project input and help in (leukemic) sample processing. We also thank Alex Q. Hoogkamer for help with analysis, the Vlietland Ziekenhuis in Schiedam for providing cord blood, and the MultiPlex Core Facility in the Center for Translational Immunology for performing Luminex experiments (Mariska van Dijk and Mariëlle Tempelman in particular). This study was funded by the Pediatric Oncology Foundation Rotterdam (SKOCR) and the Oncode Institute in Utrecht.

## Contributors

MS contributed to study design, performed the experiments, collected data, performed data analysis, and wrote the paper for this study. EMPS performed pilot work which led to this study. MLdB and CvdV designed the study and finalized this manuscript. JO, FS, MMPV, KV, and CvdV performed experiments. AQH performed RNA sequencing data analysis. SN was responsible for the Luminex experiment. All authors reviewed the final manuscript.

## Notes

### Competing Interest Statement

The authors have declared no competing interest.

