## Supplemental material for "B-Cell Precursor Acute Lymphoblastic Leukemia elicits an Interferon-α/β response in Bone Marrow-derived Mesenchymal Stroma"

### Supplementary tables

**Supplementary table 1. Characteristics of mesenchymal stromal cells derived from pediatric BCP-ALL patients**

| Mesenchymal stromal cells | Subtype ALL | Remark |
|---|---|---|
| **MSC#1** | *ETV6-RUNX1* | Relapse |
| **MSC#2** | Hyperdiploid | Relapse |
| **MSC#3** | B-other | Initial |

**Supplementary table 2. Characteristics of BCP-ALL samples**

| Leukemia (ALL#) | Subtype | Remark |
|---|---|---|
| **1** | B-other | *ETV6-RUNX1*-like |
| **2** | B-other | No fusion detected by RNAseq |
| **3** | B-other | *P2RY8-CRLF2* fusion |
| **4** | B-other | *IGH-DUX4*-rearranged |
| **5** | B-other | *IGH-EPOR*-rearranged |
| **6** | B-other | *MEF2D-BCL9* fusion |
| **7** | High hyperdiploid | |
| **8** | *ETV6-RUNX1* | |
| **9** | *ETV6-RUNX1* | |
| **10** | *ETV6-RUNX1* | |
| **11** | *ETV6-RUNX1* | |
| **12** | *ETV6-RUNX1* | |
| **13** | *ETV6-RUNX1* | |
| **14** | *ETV6-RUNX1* | |
| **15** | *ETV6-RUNX1* | |

**Supplementary table 3. Contamination percentage sorted MSC and ALL samples**

Table representing percentage of ALL cells in sorted MSC fraction and percentage MSCs in sorted ALL fraction. To avoid a bias in gene expression level due to contaminating cells, we only used sorted MSC fractions if the maximum of contaminating ALL cells was below 5% (with a few exceptions because of precious samples for which no repeated sort-experiment could be performed). Samples with contamination above 0.3% or 5.0% for ALL and MSC fractions, resp. are indicated in bold. It was very difficult to prevent contamination with MSCs in the sorts of ALL cells. Since MSCs contain more RNA than ALL (see ***Supplementary Figure 1F/G***), the maximum percentage of contaminating MSCs in ALL sorts was set to 0.3%. For 17/45 sorted samples, the percent of MSCs exceeded 0.3%. For this reason, total RNA sequencing analysis of sorted ALL cells was not included.

| **Sample** | **% ALL** | **Sample** | **% MSC** |
|---|---|---|---|
| **MSC#1 after ALL#1** | 0.80 | **ALL#1 after MSC#1** | 0.26 |
| **MSC#2 after ALL#1** | 0.82 | **ALL#1 after MSC#2** | 0.26 |
| **MSC#3 after ALL#1** | 1.56 | **ALL#1 after MSC#3** | **0.37** |
| **MSC#1 after ALL#2** | 0.46 | **ALL#2 after MSC#1** | 0.26 |
| **MSC#2 after ALL#2** | 0.60 | **ALL#2 after MSC#2** | **0.50** |
| **MSC#3 after ALL#2** | 0.69 | **ALL#2 after MSC#3** | ? |
| **MSC#1 after ALL#3** | 0.88 | **ALL#3 after MSC#1** | **0.58** |
| **MSC#2 after ALL#3** | 3.65 | **ALL#3 after MSC#2** | **0.47** |
| **MSC#3 after ALL#3** | 0.71 | **ALL#3 after MSC#3** | **0.50** |
| **MSC#1 after ALL#4** | 1.10 | **ALL#4 after MSC#1** | 0.27 |
| **MSC#2 after ALL#4** | 2.80 | **ALL#4 after MSC#2** | 0.11 |
| **MSC#3 after ALL#4** | 1.10 | **ALL#4 after MSC#3** | 0.12 |
| **MSC#1 after ALL#5** | 1.70 | **ALL#5 after MSC#1** | 0.19 |
| **MSC#2 after ALL#5** | 0.50 | **ALL#5 after MSC#2** | 0.13 |
| **MSC#3 after ALL#5** | 2.12 | **ALL#5 after MSC#3** | 0.24 |
| **MSC#1 after ALL#6** | **7.13** | **ALL#6 after MSC#1** | 0.17 |
| **MSC#2 after ALL#6** | **8.60** | **ALL#6 after MSC#2** | 0.17 |
| **MSC#3 after ALL#6** | **17.60** | **ALL#6 after MSC#3** | 0.12 |
| **MSC#1 after ALL#7** | 1.76 | **ALL#7 after MSC#1** | **0.45** |
| **MSC#2 after ALL#7** | 2.00 | **ALL#7 after MSC#2** | 0.20 |
| **MSC#3 after ALL#7** | 1.50 | **ALL#7 after MSC#3** | 0.24 |
| **MSC#1 after ALL#8** | 1.50 | **ALL#8 after MSC#1** | 0.04 |
| **MSC#2 after ALL#8** | 2.77 | **ALL#8 after MSC#2** | 0.19 |
| **MSC#3 after ALL#8** | 3.35 | **ALL#8 after MSC#3** | 0.22 |
| **MSC#1 after ALL#9** | 1.44 | **ALL#9 after MSC#1** | 0.27 |
| **MSC#2 after ALL#9** | 3.40 | **ALL#9 after MSC#2** | 0.10 |
| **MSC#3 after ALL#9** | 1.20 | **ALL#9 after MSC#3** | 0.25 |
| **MSC#1 after ALL#10** | 3.20 | **ALL#10 after MSC#1** | **0.77** |
| **MSC#2 after ALL#10** | 1.50 | **ALL#10 after MSC#2** | 0.21 |
| **MSC#3 after ALL#10** | 2.20 | **ALL#10 after MSC#3** | 0.18 |
| **MSC#1 after ALL#11** | **9.30** | **ALL#11 after MSC#1** | **0.82** |
| **MSC#2 after ALL#11** | 4.60 | **ALL#11 after MSC#2** | **0.39** |
| **MSC#3 after ALL#11** | 0.83 | **ALL#11 after MSC#3** | 0.14 |
| **MSC#1 after ALL#12** | 0.25 | **ALL#12 after MSC#1** | 0.10 |
| **MSC#2 after ALL#12** | 0.47 | **ALL#12 after MSC#2** | **0.77** |
| **MSC#3 after ALL#12** | 0.39 | **ALL#12 after MSC#3** | **0.63** |

| | | | |
|---|---|---|---|
| **MSC#1 after ALL#13** | 4.50 | **ALL#13 after MSC#1** | **0.48** |
| **MSC#2 after ALL#13** | 1.99 | **ALL#13 after MSC#2** | **0.45** |
| **MSC#3 after ALL#13** | 0.34 | **ALL#13 after MSC#3** | **0.31** |
| **MSC#1 after ALL#14** | 1.18 | **ALL#14 after MSC#1** | 0.24 |
| **MSC#2 after ALL#14** | **6.92** | **ALL#14 after MSC#2** | 0.14 |
| **MSC#3 after ALL#14** | 2.19 | **ALL#14 after MSC#3** | 0.15 |
| **MSC#1 after ALL#15** | 4.10 | **ALL#15 after MSC#1** | **0.71** |
| **MSC#2 after ALL#15** | 3.60 | **ALL#15 after MSC#2** | **0.49** |
| **MSC#3 after ALL#15** | 1.50 | **ALL#15 after MSC#3** | **0.70** |

### Supplementary Figures

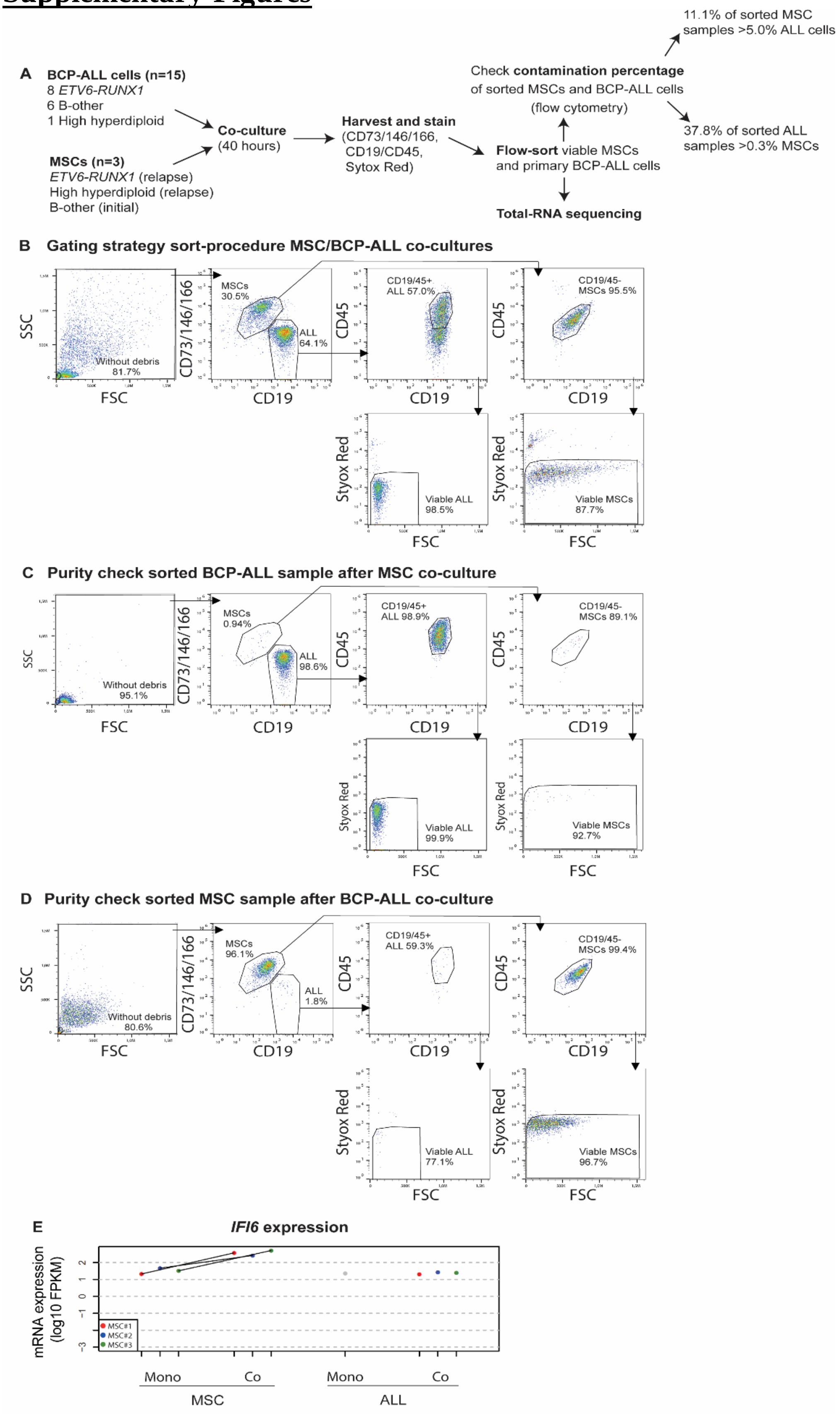

F

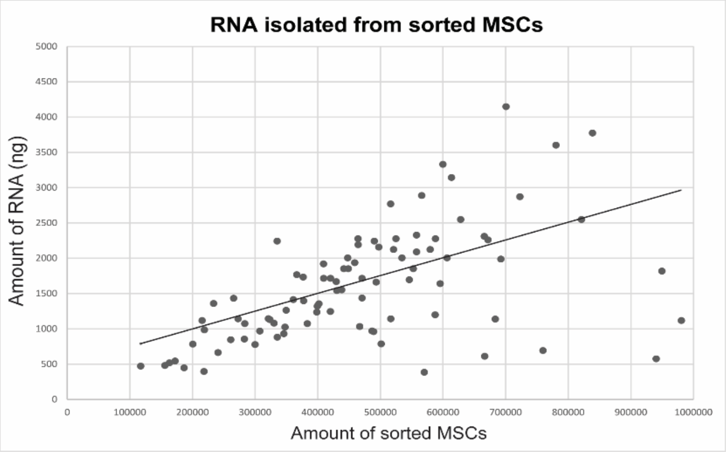


G

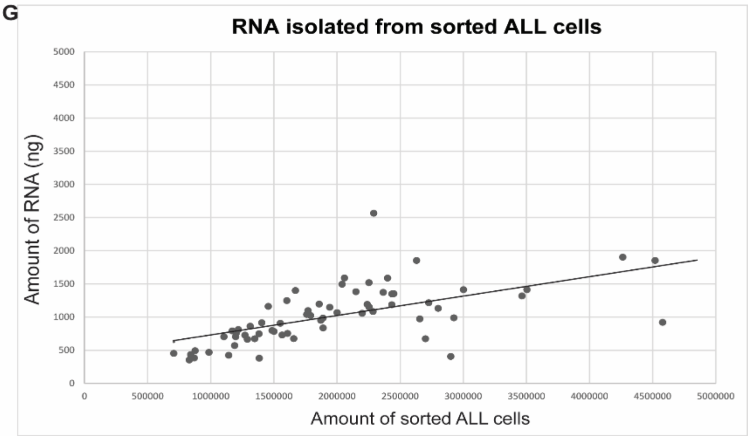


**Supplementary figure 1. Analysis of flow cytometry data and purity check of MSC and BCP-ALL mono- and co-cultures.** (A) BCP-ALL cells from 15 pediatric patients (6 B-other, 1 high hyperdiploid, 8 *ETV6-RUNX1*) were co-cultured for 40 hours with MSCs derived from 3 different origins (MSC#1: *ETV6-RUNX1*, relapse; MSC#2: high hyperdiploid, relapse; MSC#3: B-other, initial). MSCs and BCP-ALL cells were separated using FACS. Purity (contamination percentage) of sorted samples was determined by flow cytometry. RNA from sorted MSCs was used for performing total-RNA sequencing. (B) Gating strategy used for sorting MSCs and ALL#13 cells during FACS. Cell population was selected based on FSC and SSC. MSCs were selected for CD73+/CD146+/CD166+/CD19-/CD45-. BCP-ALL cells were selected for CD19+CD45+CD73-CD146-CD166-. The Sytox Red negative population represents viable cells. FSC = forward scatter, SSC = side scatter. Purity of the sorted (C) viable BCP-ALL cells and (D) MSCs after co-culture was determined by flow cytometry. (E) *IFI6* expression levels (log10-transformed FPKM) for paired MSC mono-culture and MSC after co-culture with ALL#12, ALL#12 mono-culture (indicated in grey), and ALL#12 after co-culture with MSC#1-3 indicated in red, blue, and green circle, resp. (F) Graph representing the amount of RNA isolated (y-axis) from increasing amounts (100,000-1,000,000) of sorted MSCs (x-axis). Each dot represents one sorted sample. (G) Same as (F) but for ALL cells (600,000-4,500,000).

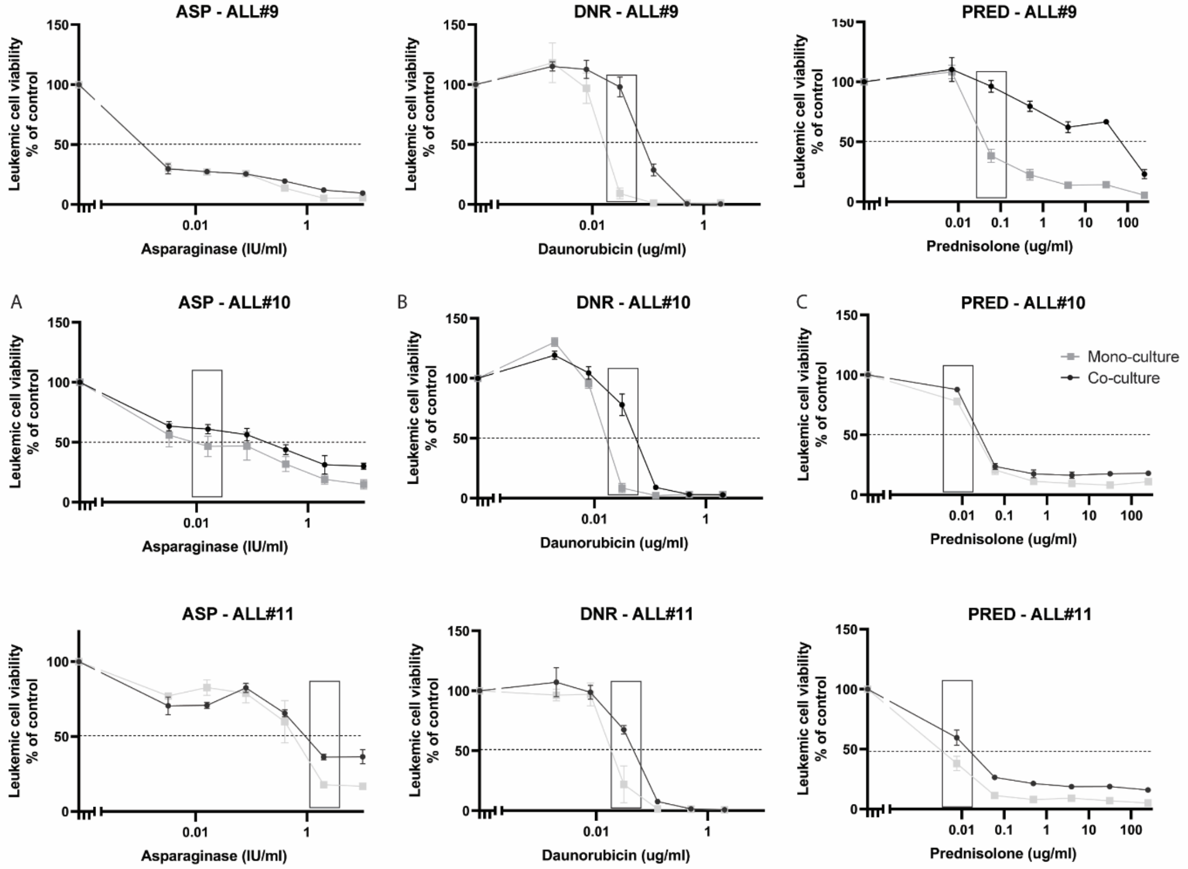


**Supplementary figure 2. MSCs protect primary BCP-ALL cells against drugs.** Graphs represent the percentage of viable primary leukemic cells (ALL#9, #10 and #11) when treated with (A) asparaginase (0.003, 0.016, 0.08, 0.4, 2.0 and 10 IU/ml), (B) daunorubicin (0.002, 0.008, 0.031, 0.125, 0.5 and 2 μg/ml) or (C) prednisolone (0.008, 0.06, 0.49, 3.9, 31.25 and 250 μg/ml) normalized to untreated control. Data points represent means of triplicate measurements ± SD. Grey squares and black circles indicate leukemic cell viability upon mono-culture and MSC co-culture, resp. Boxes indicate the drug concentration most discriminative for resistance induced by co-culturing BCP-ALL cells with MSCs. This drug concentration was used to investigate the effect of i-IFNs as shown in Figure 5. In case of ALL#9, 0.001 IU/ml asparaginase was selected as no clear effect was observed within the chosen range. Dashed line (---) indicates LC50.

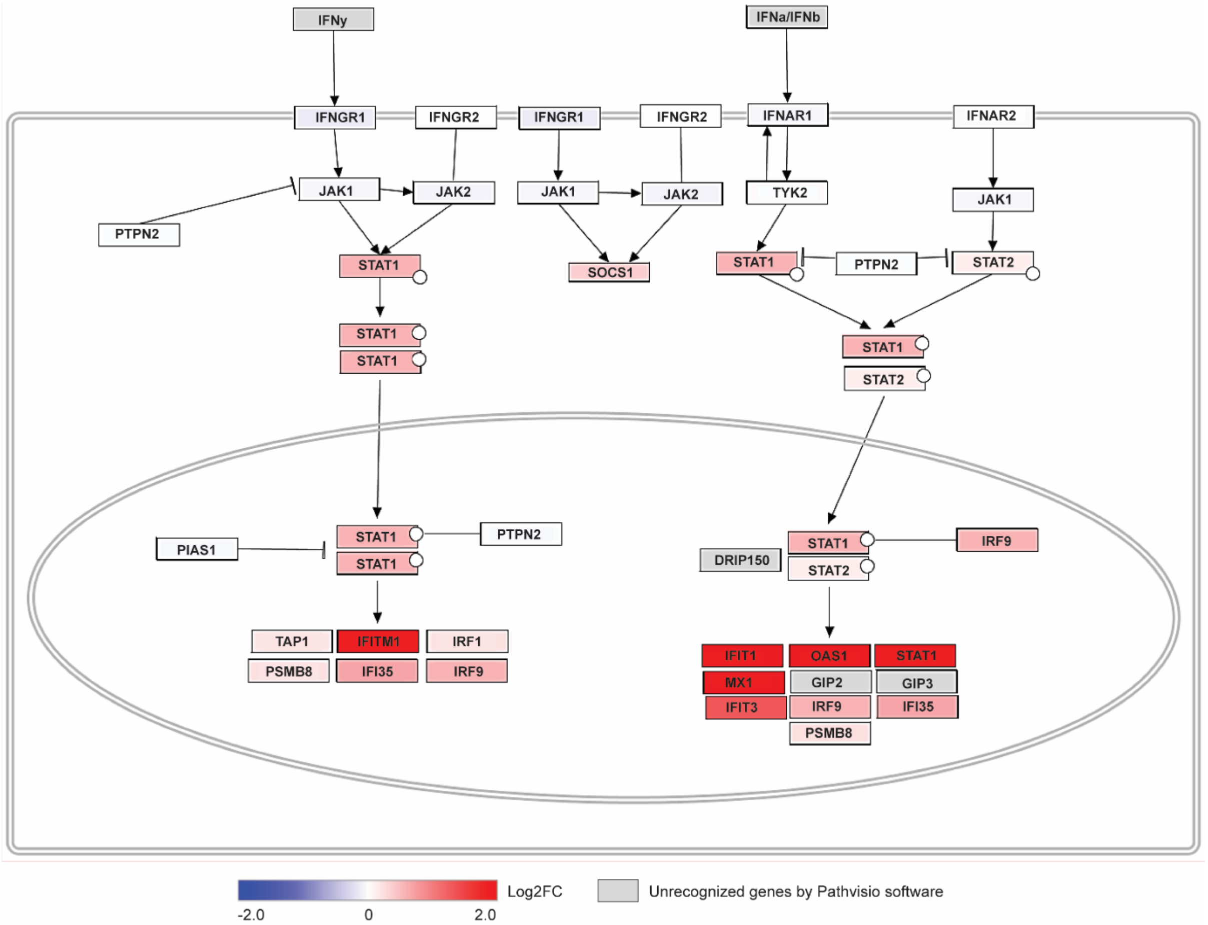


**Supplementary figure 3. Pathway analysis reveals changes in the MSCs' IFN gene profile induced by BCP-ALL cells**. Differentially expressed genes (FDR < 0.05) in MSCs (MSC#1, #2, and #3) after co-culture with primary pediatric BCP-ALL samples (n=15) analyzed by pathway analysis (Pathvisio). Blue, low expression. Red, high expression. Grey boxes indicate unrecognized genes by Pathvisio software.

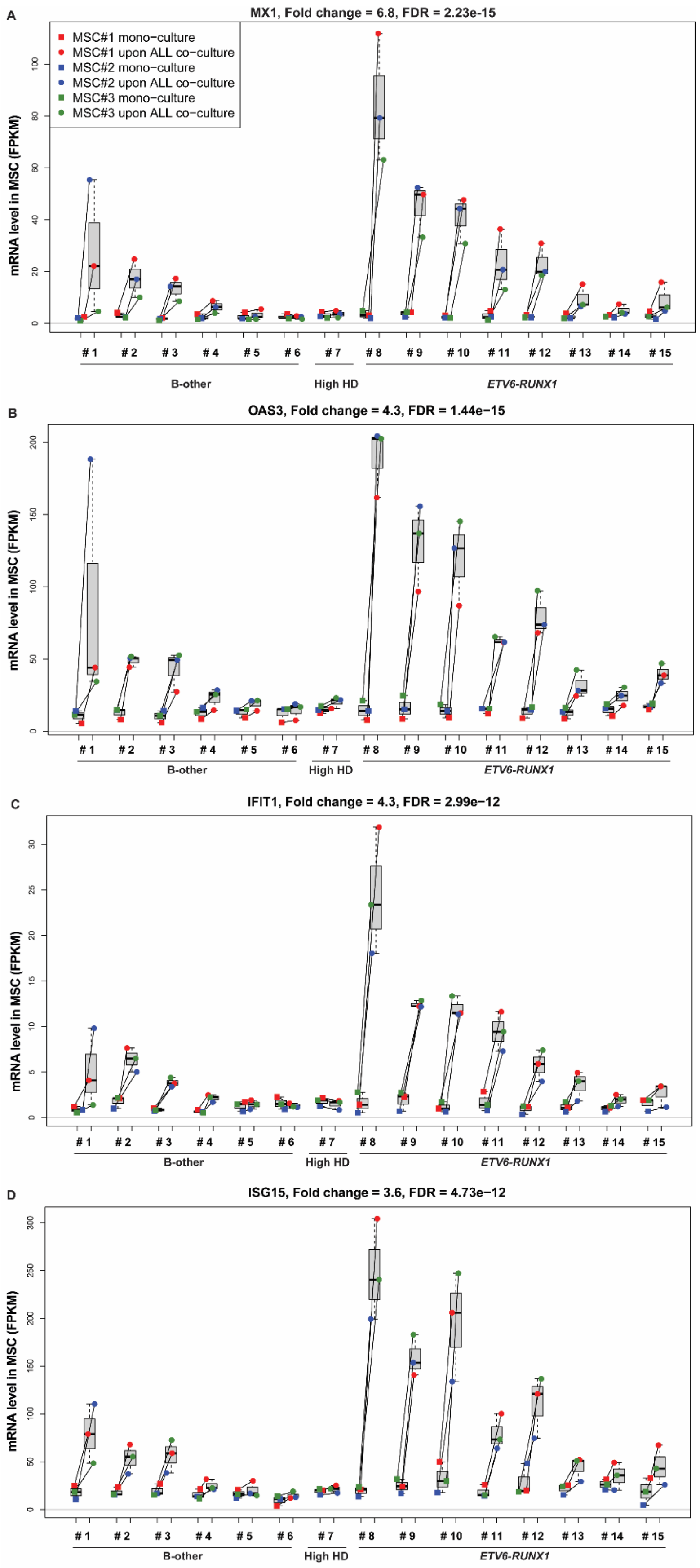

A
MX1, Fold change = 6.8, FDR = 2.23e-15
MSC#1 mono−culture
MSC#1 upon ALL co−culture
MSC#2 mono−culture
MSC#2 upon ALL co−culture
MSC#3 mono−culture
MSC#3 upon ALL co−culture
mRNA level in MSC (FPKM)
0
20
40
60
80
100
# 1
# 2
# 3
# 4
# 5
# 6
# 7
# 8
# 9
# 10
# 11
# 12
# 13
# 14
# 15
B-other
High HD
ETV6-RUNX1
B
OAS3, Fold change = 4.3, FDR = 1.44e−15
mRNA level in MSC (FPKM)
0
50
100
150
200
# 1
# 2
# 3
# 4
# 5
# 6
# 7
# 8
# 9
# 10
# 11
# 12
# 13
# 14
# 15
B-other
High HD
ETV6-RUNX1
C
IFIT1, Fold change = 4.3, FDR = 2.99e−12
mRNA level in MSC (FPKM)
0
5
10
15
20
25
30
# 1
# 2
# 3
# 4
# 5
# 6
# 7
# 8
# 9
# 10
# 11
# 12
# 13
# 14
# 15
B-other
High HD
ETV6-RUNX1
D
ISG15, Fold change = 3.6, FDR = 4.73e−12
mRNA level in MSC (FPKM)
0
50
100
150
200
250
300
# 1
# 2
# 3
# 4
# 5
# 6
# 7
# 8
# 9
# 10
# 11
# 12
# 13
# 14
# 15
B-other
High HD
ETV6-RUNX1

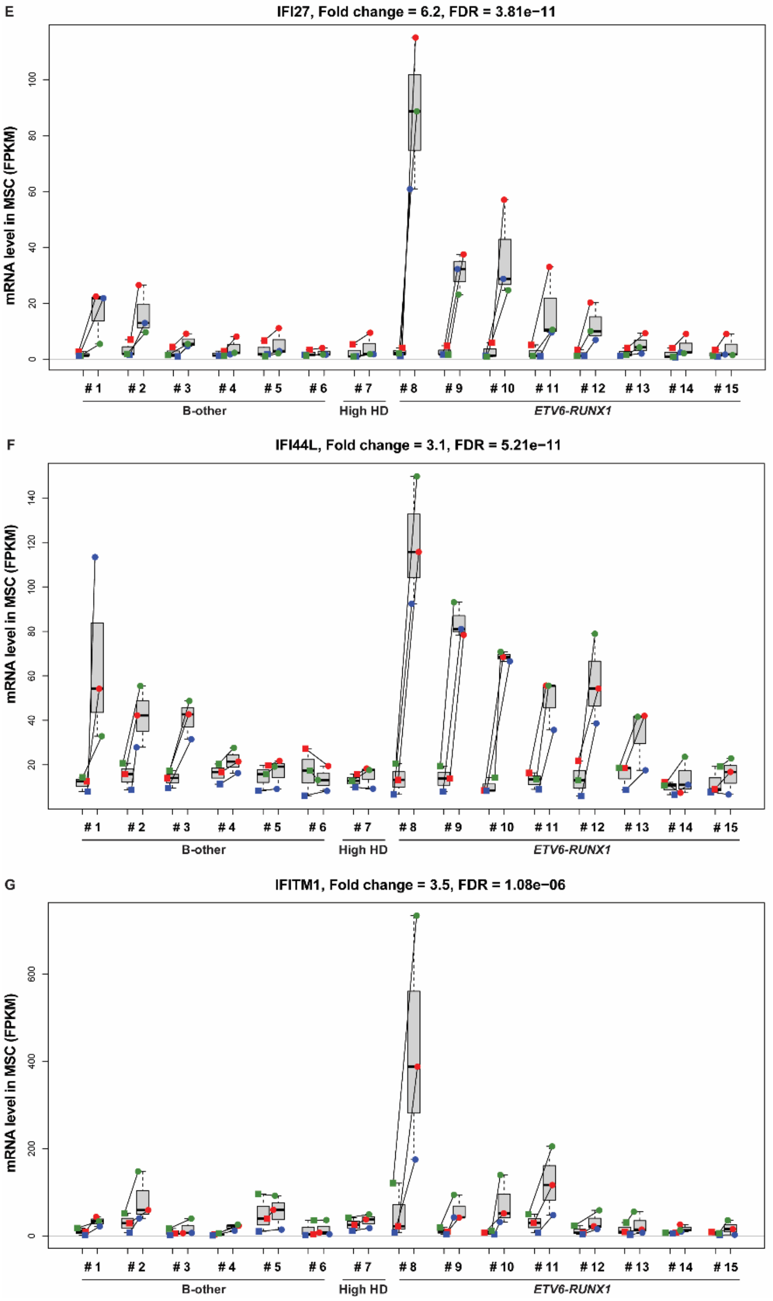


**Supplementary figure 4. Interferon-related gene signature in MSCs upon BCP-ALL co-culture.** (A) *MX1*, (B) *OAS3*, (C) *IFIT1*, (D) *ISG15*, (E) *IFI27*, (F) *IFI44L*, and (G) *IFITM1* expression levels (FPKM) for paired MSC mono-culture (squares; MSC#1-3 indicated in red, blue, and green, resp.) and MSC after co-culture with BCP-ALL cells from 15 individual patients (#1-15; circles). Boxplots represent the interquartile range, the median is depicted by a line. FDR, p-value false discovery rate. Solid grey line indicates no expression.

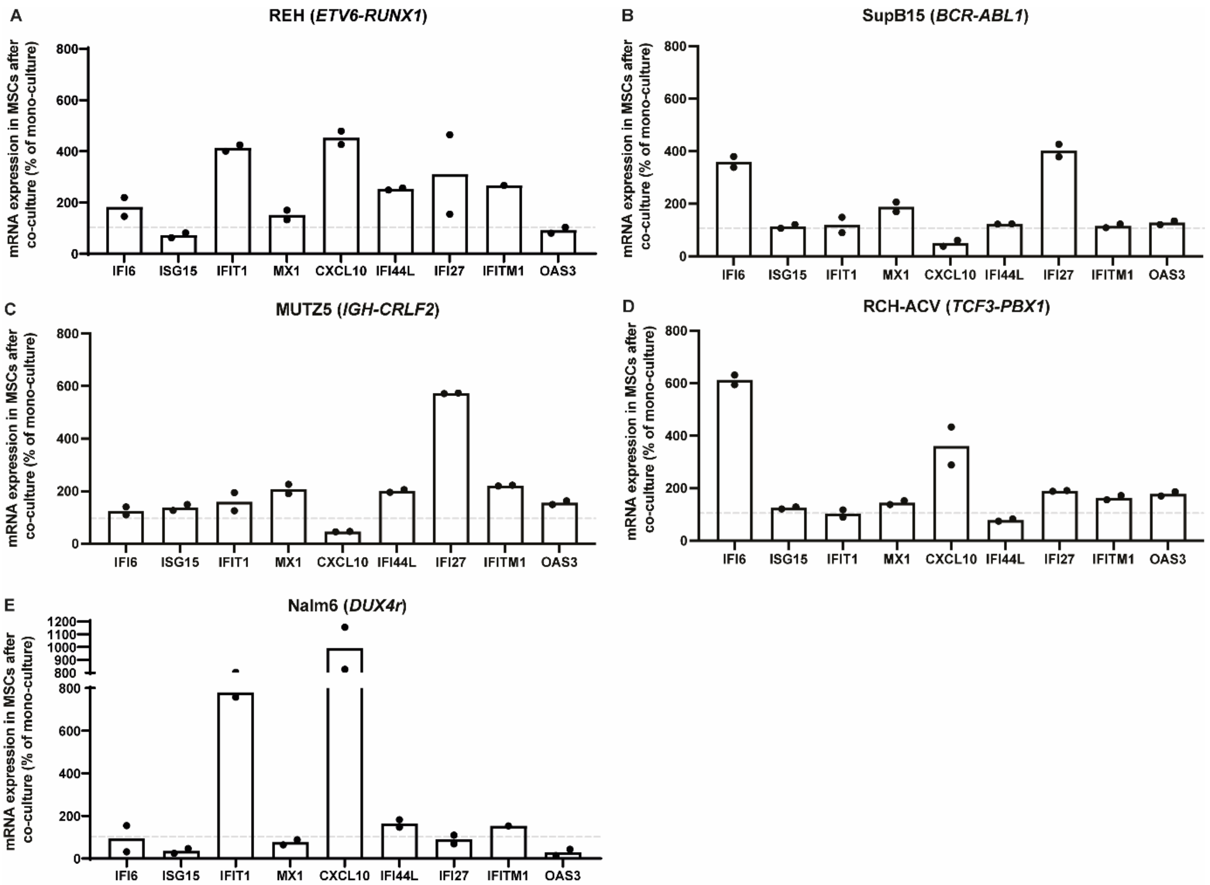


**Supplementary figure 5. IFN-gene expression levels in MSCs upon co-culture with leukemic cell lines.** mRNA expression levels of IFN-related genes in MSCs upon 40 hours co-culture with (A) REH (*ETV6-RUNX1*), (B) SupB15 (*BCR-ABL1*), (C) MUTZ5 (*CRLF2*-rearranged), (D) RCH-ACV (*TCF3-PBX1*), and (E) Nalm6 (*DUX4*-rearranged B-other) relative to MSC mono-culture. Bars are means of duplicate measurements for 1 experiment. Dashed line (---) indicates mRNA expression levels of MSC after mono-culture, set to 100%.

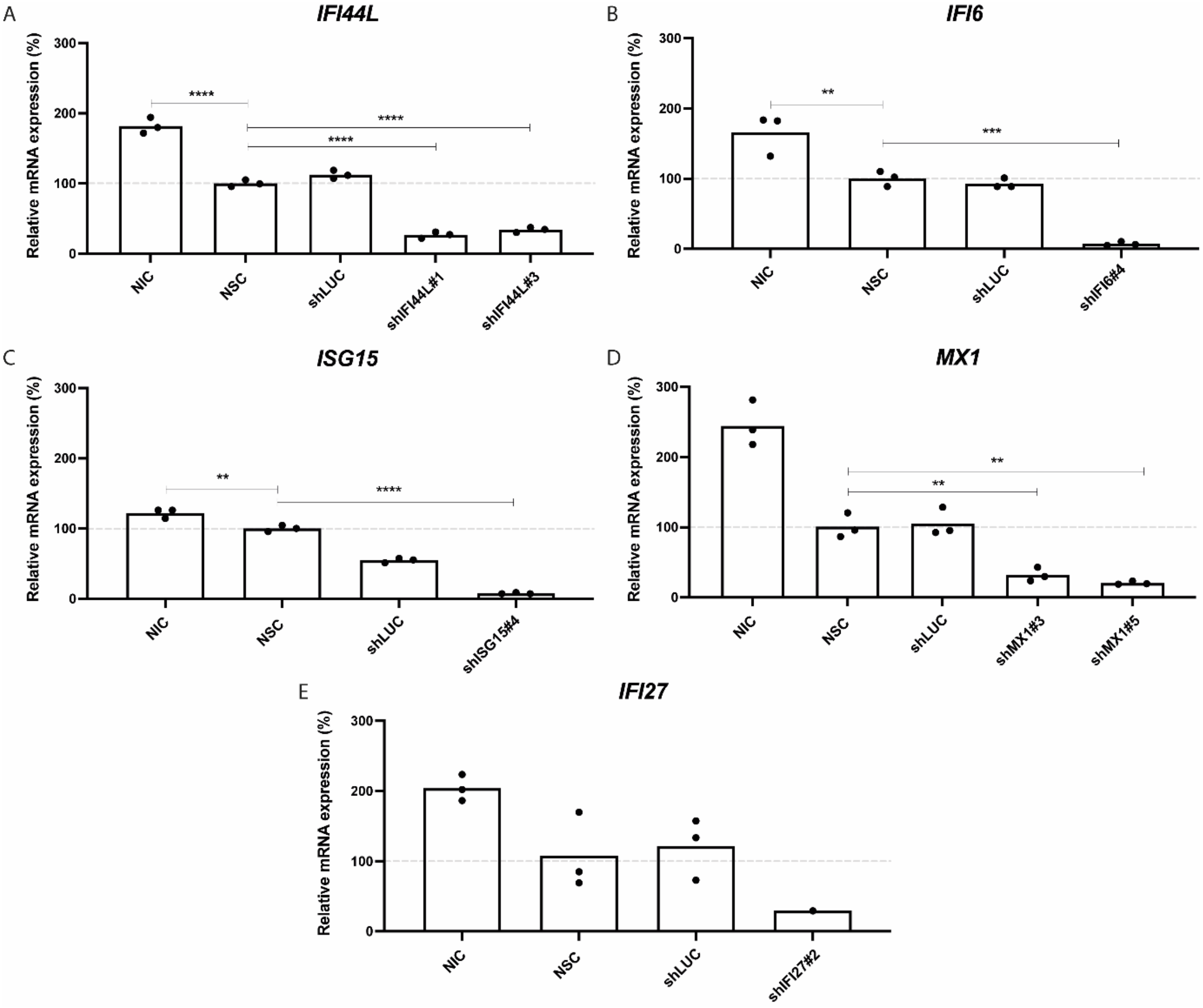


**Supplementary figure 6. Knockdown efficiency of IFN-related genes in MSCs.** A representative experiment showing relative mRNA expression levels of (A) *IFI44L*, (B) *IFI6*, (C) *ISG15, (D) MX1*, and *(E) IFI27* in MSCs (MSC#2) after lentiviral silencing. Bars represent means of technical triplicates for one independent experiment. mRNA expression was normalized to the non-silencing control (NSC). Dashed line (---) indicates mRNA expression levels of MSCs treated with shNSC, set to 100%. NIC = Non-Infected Control, shLUC = short-hairpin-Luciferase.
